# Emergence of *β* and *γ* networks following multisensory training

**DOI:** 10.1101/560235

**Authors:** Daria La Rocca, Philippe Ciuciu, Denis Alexander Engemann, Virginie van Wassenhove

## Abstract

Our perceptual reality relies on inferences about the causal structure of the world given by multiple sensory inputs. In ecological settings, multisensory events that cohere in time and space benefit inferential processes: hearing and seeing a speaker enhances speech comprehension, and the acoustic changes of flapping wings naturally pace the motion of a flock of birds. Here, we asked how a few minutes of (multi)sensory training could shape cortical interactions in a subsequent perceptual task, and investigated oscillatory activity and functional connectivity as a function of sensory history in training. Human participants performed a visual motion coherence discrimination task while being recorded with magnetoencephalography (MEG). Three groups of participants performed the same task with visual stimuli only, while listening to acoustic textures temporally comodulated with the strength of visual motion coherence, or with auditory noise uncorrelated with visual motion. The functional connectivity patterns before and after training were contrasted to resting-state networks to assess the variability of common task-relevant networks, and the emergence of new functional interactions following training. One main finding is the emergence of a large-scale synchronization in the high *γ* (gamma: 60*−*120*Hz*) and *β* (beta:15*−*30*Hz*) bands for individuals who underwent comodulated multisensory training. The post-training network involved prefrontal, parietal, and visual cortices. Our results suggest that the integration of evidence and decision-making strategies become more efficient following congruent multisensory training through plasticity in network routing and oscillatory regimes.

## 1. Introduction

The brain can infer the causal structure of its surroundings by integrating multisensory signals originating from the same physical source while segregating those originating from different causes [Par+12; PE16; Der+16; KS15; RN15]. The resolution of this causal inference problem weighs in the reliability and the degree of correspondence between multisensory inputs [Roa+06; Spe11; Mad+15]. In ecological settings, the temporal comodulation of sensory signals helps perceptual scene analysis: for instance, an interlocutor’s mouth movements are temporally coherent with the envelope of the acoustic speech signals providing the listener with strong binding cues for predictive inferences [GS00; VW+05; Sch+08b; Nah+15; Was13; Mad+15]. Temporally congruent signals enhance the detectability [Bur+08; Mad+15] and the identification [KVW12; Zil+14] of events, whereas temporally incongruent signals hinder their identification [KVW12; Mad+15]. Herein, we explored the cortical mechanisms by which the internalized temporal structure of coherent multisensory events may subsequently regulate visual processing.

Using magnetoencephalography (MEG), we characterized the impact of uni- and multi-sensory training history on human brain activity when participants (*N* = 36) performed a visual motion coherence task (Figure 1.A). The task consisted in reporting the color of the most coherent cloud of dots amongst two intermixed red and green clouds of moving dots. After initially performing the task with visual stimuli only (PRE), participants were split into three experimental groups for specific training: one group performed the task with visual stimuli only (V), another one with acoustic textures spectro-temporally congruent with the most coherent visual cloud of the two (AV) and a third one, with distracting auditory noise uncorrelated with any of the two visual clouds (CTRL). After performing the training for 20 minutes, all participants were again tested with visual stimuli only (POST). Behaviorally, all participants improved their perceptual discrimination with the AV group showing the largest perceptual benefits and the initial analysis of the MEG evoked activity suggested the implication of a large-scale network following training [Zil+14].

**Figure 1:**
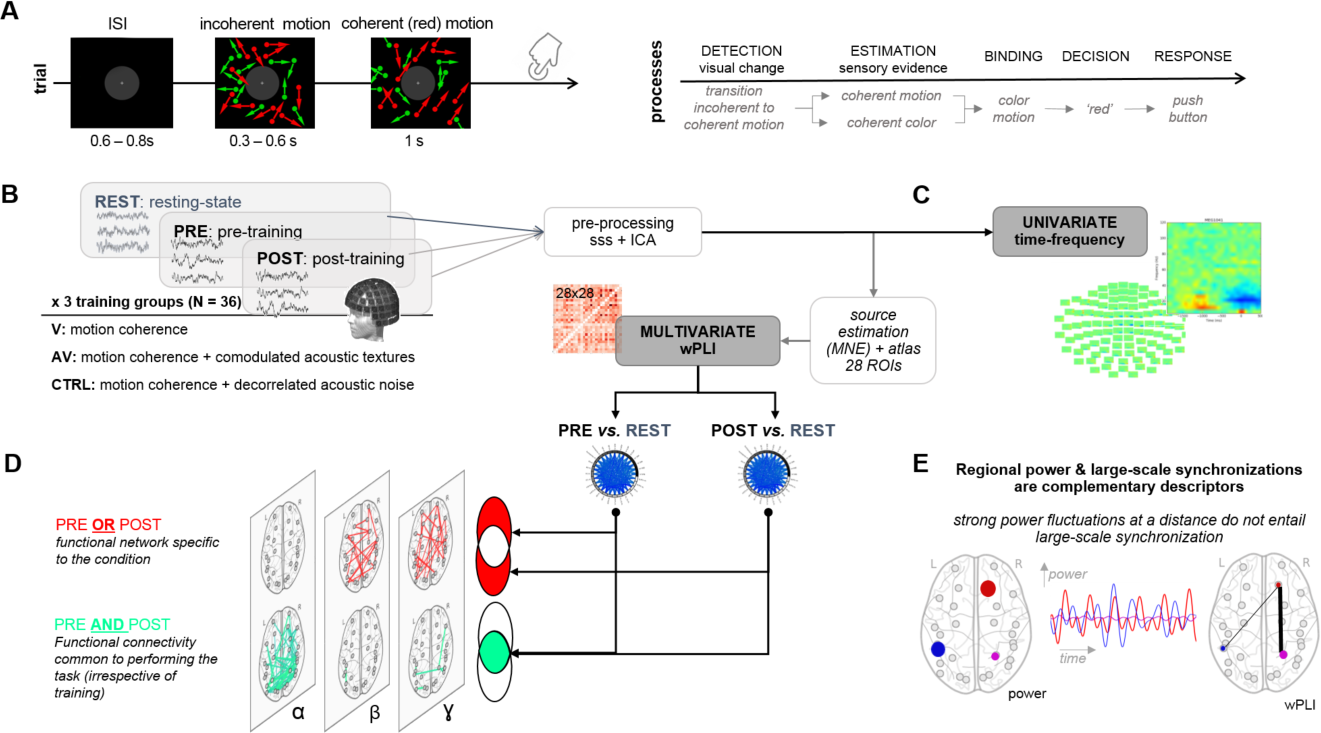
Experimental procedure and Methods. (**A**) Left panel: each trial started with the presentation of a fixation cross lasting 0.6 to 0.8 s followed by the presentation of two intermixed clouds of dots moving incoherently. One cloud was red, the other, green. After a variable delay (0.3 *−* 0.6 s) of incoherent motion, one of the clouds (here, red) moved coherently for 1 s while the other remained fully incoherent (here, green). Seven possible strength of motion coherence were tested; the direction and color were randomized across trials. Participants selected which of the red or green cloud was most coherent (videos S1 and S2 in [Zil+14]). Right panel: schematic operationalization of the motion coherence discrimination task entailing the integration of motion and color for decision-making. (**B**) MEG recordings were collected from 36 participants, who performed the task described in (A) in the PRE and POST blocks. Between the PRE and POST, participants were split in three experimental training groups who performed the task visually (V), with acoustic textures congruent with the most coherent cloud of dot (AV), or with auditory noise uncorrelated with the visual stimuli (CTRL). All new analyses were carried out on the PRE and POST blocks, when all participants were performing the visual task described in (**A**). All participants improved their behavioral scores in POST as compared to PRE blocks: the AV group showed the largest perceptual benefit followed by the V and the CTRL group [Zil+14]. Following preprocessing and source estimations, (**C**) univariate time-frequency analysis and (**D**) multivariate functional connectivity analyses were performed to provide (**E**) complementary insights on the oscillatory mechanisms implicated in the effect of (multi)sensory training history in unisensory processing.

In the present work, we re-analyzed data collected in [Zil+14] and characterized the oscillatory networks following different perceptual histories by assessing the changes of brain activity between PRE and POST blocks, when all participants performed the task with visual stimuli only (Figure 1.B). Although we did not focus on the effects of feed-forward integration of multisensory features, or selective attention on brain activity, our analytical approach built on seminal work suggesting the implication of distinct neural oscillatory coupling within large-scale networks [Sen+08; KS18], [Hip+11]). The dynamic regimes mediating the binding of multisensory information across brain regions have started being characterized [Lak+08; Sen+08; KS18; Att+14], yet little is known regarding the oscillatory networks which may actively contribute to supramodal or multisenory object representations [Was13; Zil+14; Biz+16]. Given this experimental approach, we expected a strong implication of top-down oscillatory components and networks specifically in the best learning group (AV) implicating *β* oscillations, which is a critical spectral signature of efficiency in feedback and sensorimotor decision-making [EF10] as well as a marker of integrative functions [Sie+11].

To characterize the different oscillatory networks, we estimated oscillatory activity within, and across, experimental groups using univariate time-frequency analyses (Figure 1.C) and large-scale functional connectivity (FC) measures based on the weighted phase lag index (wPLI) [Vin+11] (Figure 1.D). We investigated a large network including prefrontal, parietal, occipital and temporal cortices with regions orthogonally selected for their functional relevance in the task (cf [Zil+14], see Methods). Among regions of interest were the ventro-lateral prefrontal cortex (vlPFC), a massive site of convergence for visual, auditory and multisensory information processing [Rom07; RH12], whose neurons selectively respond to the color of visual objects [Rom12] and low-level abstraction [Wut+18]; the intra-parietal sulcus (IPS), which plays a central role in multisensory processing [And97; BM11; Pas+10] and visual motion area (MT), sensitive to perceptual changes in this task [Zil+14]. Both IPS and MT are known to interact in the *β* range during perceptual decision-making [Don+09]. We first started by exploring the modulation of local oscillatory activity during visual motion discrimination [Sie+06; Sie+11], and followed up with the exploration of changes in functional connectivity as a function of sensory history in training.

## 2. Materials and methods

### 2.1 Participants

36 healthy human participants were recruited for the study (age range: 18 to 28 y.o.; mean age: 22.1 *±*2.2 s.d.; 3 groups of 12 participants each: V: 4 females; AV: 6 females; AVr: 6 females). All participants were right-handed, had normal hearing and normal or corrected-to-normal vision. Before the experiment, all participants provided a written informed consent in accordance with the Declaration of Helsinki (2008) and the local Ethics Committee on Human Research at NeuroSpin (Gif-sur-Yvette, France). Prior to the MEG acquisition, participants were randomly split into 3 experimental groups (V, AV, and CTRL) as detailed below.

### 2.2 Task

The MEG experiment consisted of interleaved MEG blocks alternating between rest and task. The task blocks included: a 12 minutes pre-training block (PRE) consisting of the visual coherence discrimination task; a 20 minutes training (4 successive blocks of 5 minutes each) on the same task using purely visual stimuli (V group), congruent audiovisual stimuli (AV group) or incongruent audiovisual stimuli (CTRL group); a 12 minutes post-training block (POST) consisting of the same visual coherence discrimination task as in PRE. Thus, the PRE and POST blocks consisted of the same visual *only* coherence discrimination task for all three experimental groups and using the exact set of visual stimuli. Only the training was either visual or audiovisual. The task requirements in PRE, training, and in POST were otherwise identical in all runs: two clouds of colored dots were intermixed on the screen and participants had to tell which of the red or green cloud of dots was the most coherent. In PRE and POST, participants also rated their confidence on a scale of 1 to 5 after they provided their main response regarding the color of the most coherent cloud of dots.

In PRE and POST, the initial and final motion coherence discrimination threshold of each participant was assessed by testing seven strength of visual motion coherence (15%, 25%, 35%, 45%, 55%, 75% and 95%). 28 trials for each strength of visual motion coherence were collected in PRE and in POST for a total of 196 trials in each block. In the training (4 blocks, 5 min each), four visual coherence levels were tested corresponding to *±* 10% and *±*20% of an individuals discrimination threshold computed in PRE (see [Zil+14] for more details). 28 trials for each strength of visual motion coherence were presented for a total of 112 trials in a given training block. These data were not considered in this study as our main question focused on contrasting brain activity to identical experimental conditions given a different training history. Further experimental details can be found in [Zil+14].

The first resting block occurring prior to any task or training will be thereafter referred to as REST and was used as baseline for functional connectivity analysis.

### 2.3 Stimuli

Visual stimuli consisted of intermixed red and green clouds of dots (Figure 1.A) calibrated to isoluminance using heterochromatic flicker photometry on a per individual basis prior to MEG data acquisition. A white fixation cross was at the center of a 4° gray mask disk and dots were presented within an annulus of 4° to 15° of visual angle. Dots had a radius of 0.2°. The motion flow was 16.7 dots per deg^2^*×* s with a speed of 10°/s and its direction confined within an angle of 45° − 90° around the azimuth. 50% of the trials were upward coherent motion and the remaining 50% of the trials were downward coherent motion. The color and the direction of the most coherent cloud o dots were thus pseudo-randomized across trials and both color and direction were orthogonal to the task goal.

The V group underwent training using visual only stimuli. The AV group underwent training using temporal comodulated audiovisual associations comparable to those used in sensory substitution devices such as the vOICe [Mei92] and the EyeMusic [LT+12], with intuitive perceptual associations between sensory modalities [MO87; Mae+04]. Here, we used parametric sounds or acoustic textures (cf [Ove+10] with sampling frequency = 44.1 kHz, frequency range: 0.2 to 5 kHz) which enabled to pair each visual dot with a linear frequency-modulated acoustic sweep whose slope depended on the direction taken by the visual dot (see [Zil+14] for more details). The maximal slope was 16 octaves/s corresponding to motion directions of 82.9° − 90°. A visual dot moving upwards was associated with an upward acoustic ramp, whereas a downward moving dot was associated with a descending acoustic ramp. The duration of a ramp was also identical to the life-time of a visual dot. The CTRL group underwent training with acoustic noise of the same duration and amplitude as the acoustic textures used for the AV group. Crucially, the acoustic noise was thus fully uncorrelated with the visual coherent motion and served as a control regarding the specificity of audiovisual associations in this task.

In the task and for all experimental groups, a given trial started with a variable duration (0.3 to 0.6*s*) mixing both red and green clouds of dots being fully incoherent (0% of coherent motion). Then, one cloud of dots became more coherent than the other for a duration of one second. In PRE and POST, the coherence level taken by the most coherent cloud was one of seven possible values described in the Task section. During training, the coherence level taken by the most coherent cloud took one of four values described in the Task section. Inter-trials intervals (ITI) varied from 0.6 to 0.8*s*. Samples of the video trials can be experienced (Movies S1 and S2 in [Zil+14]).

### 2.4 MEG and MRI data acquisition

Electromagnetic brain activity was recorded in a magnetically shielded room using a 306 MEG system (Neuromag Elekta LTD, Helsinki). MEG signals were sampled at 2 kHz and band-pass filtered between 0.03-600 Hz. Four head position coils (HPI) were used to measure the head position of participants before each block; three fiducial markers (nasion and pre-auricular points) were used during digitization as a reference for coregistration of anatomical MRI (aMRI) immediately following MEG acquisition. Electrooculograms (EOG) and electrocardiogram (ECG) were recorded simultaneously with MEG. Five minutes of empty room recordings were acquired before each block for the computation of the noise covariance matrix.

The T1 weighted aMRI was recorded using a 3-T Siemens Trio MRI scanner. Parameters of the sequence were: field-of-view: 256*×*256*×*176 mm^3^ (transversal orientation), voxel size: 1.0 *×* 1.0 *×* 1.1 mm; acquisition time: 466 s; echo time TE = 2.98 ms, inversion time TI = 900 ms, repetition time

TR = 2300 ms and flip angle (FA): 9°. For each participant, cortical reconstruction and volumetric segmentation of T1 weighted aMRI was performed using FreeSurfer^1^. Once cortical models were complete, deformable procedures were executed using the MNE software [Gra+14] to register source estimates of each individual onto the FreeSurfer average brain for group analysis.

### 2.5 MEG preprocessing

The analysis of the MEG data was carried out using the MNE-python toolbox [Gra+13]. After applying an anti-aliasing FIR filter (low-pass cut-off frequency at 130 Hz), MEG data were down-sampled to 400 Hz, and preprocessed (Figure 1.B) to remove external and internal interferences, in accordance with accepted guidelines for MEG research [Gro+13]. Signal

Space Separation (SSS) was applied with MaxFilter to remove exogenous artifacts and noisy sensors [TS06]. Ocular and cardiac artifacts (eye blinks and heart beats) were removed using Independent Component Analysis (ICA) on raw signals. ICA were fitted to raw MEG signals, and sources matching the ECG and EOG were automatically found and removed before signals reconstruction following the procedure described in [Gra+14]^2^.

### 2.6 Univariate time-frequency analysis in sensor space

The oscillatory activity in *α*, *β* and *γ* ranges was established using a univariate time-frequency analytical approach in sensor space(Figure 1.C). Oscillatory activity in the post-stimulus (POST) period was contrasted with pre-stimulus (PRE) period. Due to the variability in the duration of the initial incoherence period of the stimuli as well as the natural variability in reaction times (RTs), data were locked and epoched according to three different types of events. A first epoching ranged from *−*600 ms to +900 ms *post-incoherent* motion onset (initial stimulus display); the second epoching ranged from 0 ms to +1500 ms *post-coherent* motion onset; the third epoching ranged from *−*1000 to +500 ms around the *button press* (RT). Importantly, the 600 ms interval preceding the incoherence onset was used as baseline for all sets of epochs.

For each set of epochs, a group-level non-parametric spatio-temporal cluster analysis was computed on single-trial time-frequency transform of the signals obtained with Morlet complex-valued wavelets and averaged in each frequency band of interest. The number of cycles in the Morlet wavelet was defined for each frequency (*f*) as *f/*2. To assess the statistical significance of the obtained clusters we randomly flipped *r* = 10^4^ times the sign of the time-frequency transformed data, and cluster-level correction for multiple comparisons was based on the maximum statistic method [MO07]. The spatio-temporal clustering was used to identify the sensors showing event-related activity and latencies in PRE and in POST. One of these sensors was used for the subsequent univariate analysis. This ensured that any inference made on a particular frequency band was first determined independently of an ad-hoc working hypothesis motivating the subsequent contrasts. Specifically, time-frequency cluster analyses was used in PRE to perform group-level statistics (*N* = 36, low *vs.* high MC, and correct *vs.* incorrect trials). Moreover, a statistical contrast was performed between POST and PRE blocks, pooling all participants together (*N* = 36) as well as considering each group separately (*N* = 12). Statistical significance for all these contrasts was assessed using random permutations as discussed above.

To test the modulation of oscillatory activity during task execution, we used a general linear model (GLM) on the power of the oscillatory activity regimes averaged on a single trial basis and over specific time intervals defined by significant latencies found in the cluster analysis. A non-parametric approach to GLM based on random permutations was employed to obtain a robust and unbiased linear regression [AL99; DR17]. A GLM was employed to test linear regression according to the equation *y* = **w^T^x** + *ε*. Here *y ∈* ℝ was the MEG mean power in a significant cluster of sensors; **x** was the vector [1, x_1_, x_2_, …, x_p−1_]^*T*^ ∈ ℝ^p^ containing *p* regressor variables. To find the best fitting model, we tested different combinations of regressors including motion coherence, reaction times, correctness, confidence ratings and their interactions. In particular, each regressor was first tested in an independent linear model, and significant explanatory variables were subsequently tested in the same model, together with their interactions in order to identify possible driving effects. **w** contained the *p* regression coefficients including the constant term, and *ϵ* was the error term. Iteratively reweighted least squares was used to obtain an estimate of **w** and a value of the Wald statistic *w_ref_*. A non-parametric approach based on random permutations was used to obtain robust and unbiased significance levels and confidence intervals. Specifically, to test the significance of each estimated regression coefficient *w_i_*, *r* = 10,000 random permutations of the corresponding regressor variable *x_i_* were generated, yielding a distribution of Wald statistics *w^∗^* for each partial regression coefficient under the null hypothesis *H*_0_ : *w_i_* = 0. For each estimated coefficient, the p-value was calculated as the proportion of *w^∗^* grater then or equal to *w_ref_*, in absolute value. Permutation inference for the GLM in common neuroimaging applications has been proposed as a non-parametric test to relax assumptions on data distributions [Win+14]. The 36 participants were pooled together in PRE (*N* = 36) whereas group-specific analyses (*n* = 12) were performed on POST data to study the effects of (multi)sensory training. This analysis was carried out for the three sets of epochs locked to the three different events (incoherence onset, coherence onset, response).

### 2.7 MRI-MEG coregistration and source reconstruction

The coregistration of MEG data with the individual aMRI was carried out by realigning the digitized fiducial points with the markers in MRI slices, using MRILAB (Neuromag-Elekta LTD, Helsinki) and *mne analyze* tools within MNE ([Gra+14]). Individual forward solutions for all source reconstructions located on the cortical sheet were computed using a 3-layers boundary element model constrained by the individual aMRI. Cortical surfaces were extracted with FreeSurfer and decimated to about 5,120 vertices per hemisphere with 4.9 mm spacing. The forward solution, noise and source covariance matrices were used to calculate the noise-normalized dynamic statistical parametric mapping (dSPM) ([Dal+00]) inverse operator (depth = 0.8). The inverse solution was obtained using a loose orientation constraint on the transverse component of the source covariance matrix (loose = 0.4). The estimates of the reconstructed dSPM time series were interpolated onto the FreeSurfer average brain for group-level source space analysis. Only the radial components of the estimated currents were considered for further analysis.

After source estimation, 28 regions of interest (ROIs) covering both hemispheres were identified on each individual cortex by using FreeSurfer cortical parcellations^3^. The obtained ROIs comprised regions shown to be involved in multisensory processing, perceptual decision making and motion discrimination ([Zil+14]), together with control regions, namely: frontopolar regions (FP), frontal eye field (FEF), ventro-lateral prefrontal cortex (vlPFC), pre-motor cortex and supplementary motor region (BA6), primary motor cortex (PMC), intra-parietal sulcus (IPS), inferior temporal cortex (ITC), auditory cortex (AUD), superior temporal sulcus (aSTS, mSTS and pSTS), middle temporal visual area (MT), visual area V4, and primary and secondary visual cortices (V1-V2). The average activities over all the vertices within each of these cortical regions (labels) were used for the subsequent functional connectivity analysis.

### 2.8 Functional connectivity analysis

#### 2.8.1 Adjacency matrices

Functional interaction between brain regions was assessed by evaluating the similarity of brain activity across remote brain areas, namely functional connectivity (FC) (Figure 1.D). Several studies have compared a subset of FC methods with respect to their ability to correctly detect the presence of simulated connectivity schemes in a multivariate data set [AA+06]. The outcomes showed that the performance of the measures depended on the characteristics of the dataset and the methods. No single method out-performed the others in all cases. A practical and reasonable approach thus consisted in predetermining the FC method according to the plausible working hypotheses of the experimental study. To characterize FC in the absence of *a priori* knowledge about its nature and the generating model, non-parametric measures could first be used.

The notion of phase coupling derives from the study of oscillatory nonlinear dynamical systems. Based on this notion, Phase Lag Index (PLI) [Sta+07] aims at quantifying in a statistical sense the phase delay between such systems from experimental data [Saz+09] according to the following formula:

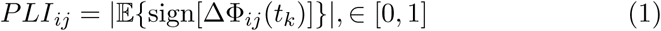

where Δ_*ij*_Φ(*t_k_*) = Φ_*i*_(*t_k_*) *−* Φ_*j*_(*t_k_*) quantifies the instantaneous phase difference between two source reconstructed time series *s_i_*(*t*) and *s_j_*(*t*) at time point *t* = *t_k_*. In Eq. (1), the expectation is typically replaced by the empirical mean over consecutive time points. PLI was shown to be robust with respect to instantaneous linear mixing effects which may lead to the detection of spurious functional couplings not caused by brain interactions (instantaneous linear mixing effects) [Sta+07].

Moreover, PLI has the advantage of not being influenced by the magnitude of phase delays. Weighted PLI (wPLI) also solves the problem of discontinuity around zero [Vin+11], by using the magnitude of the imaginary part of the cross-spectrum as weights. To measure pairwise interactions between the extracted cortical labels, we used the definition of wPLI in the frequency domain, exploiting the phase of the Fourier-based cross-spectrum *S_i,j_*(*f*) of two time series *s_i_*(*t*) and *s_j_*(*t*):

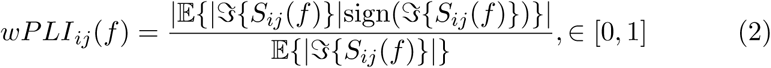

where 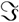 stands for the imaginary part, and the expectations were replaced by their empirical estimates averaged over epochs. Here, *f* usually spans a specific frequency band such as oscillatory regimes (*α*, *β* or *γ*). Therefore, each FC observation consisted of a symmetric adjacency matrix of size 28 × 28. 10 instances of FC were obtained for each participant and each block performing a partition of epochs into 10 non-overlapping subsets. In order to ensure the balance of the number of epochs used to obtain each FC instance for the different participants, the total number of epochs was set to the minimum observed across participants.

#### 2.8.2 Statistical analysis of FC

A widely employed approach to extract the FC network of interest from an adjacency matrix consists in applying a threshold to the strength of the estimated connections (Figure 4.A). The threshold is obtained according to a suitable criterion [DVF+14]. The resulting FC patterns correspond to the strongest connections, which do not necessarily reflect the most significant differences between experimental conditions. Additionally, while such approach is particularly suitable for graph-theoretic network analysis, it does not allow direct quantitative comparisons, owing to the variability of significant connections.

Here, the goal was to separately investigate changes that were task-dependent (i.e. significant changes in the contrast PRE or POST *vs.* REST) and cortical interactions driven by (multi)sensory training. Hence, the comparison between FC estimates obtained for the three experimental groups was addressed using a different approach. First, adjacency matrices were separately averaged over each frequency band of interest, each block (REST, PRE and POST) and each participant (Figure 4.A). Second, for each frequency band and each experimental group (AV, V, and CTRL), the task-relevant networks were extracted by performing a group-level permutation t-test between FC estimated in REST and FC estimated in task blocks (PRE, POST) (Figure 4.B). Third, considering only the subset of task-related connections common to PRE and POST blocks (i.e. the connections significantly changing both in PRE and POST as compared to REST), the variability driven by the training (POST *vs.* PRE) was evaluated using a permutation t-test (Figure 4.C, top).

All statistical tests were corrected for multiple comparisons using the maximum statistic method [MO07]. Finally, the reorganization of FC in POST was addressed by highlighting the emergence of new task-relevant FC in POST, which were not observed in PRE (Figure 4.C, bottom). Importantly, this approach considered the FC at REST as the baseline for all other FC analyses. This allowed to better disentangle the different FC patterns and their changes between PRE and POST. Hence a linear correlation analysis based on Pearson’s correlation coefficient was performed between the average increase of post-specific interactions from PRE to POST and the corresponding increase of confidence ratings, for each frequency band and each training group separately.

#### 2.8.3 Topological analysis of FC

A complementary and conventional topological analysis of FC networks was also carried out [BS09]. Specifically, the networks with density threshold given by 3*/N_rois_*, where *N_rois_* is the number of regions [DVF+17], were first extracted for each participant, each block and each frequency band. Weighted node degree *D_i_*, a topological property which is a conceptually simple measure of centrality of a node *i* within a network, was then computed for each label in the extracted networks, according to the formula:

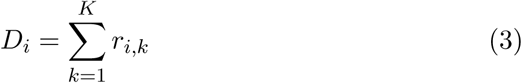

where *K* is the number of nodes in the network (cortical labels), and *r_i,k_* is the estimated FC value between nodes *i* and *k*. Permutation t-test were performed to evaluate the differences of node degree values between REST and task blocks (i.e. PRE or POST) as well as between PRE and POST. Again, the maximum statistic method was used to correct the statistical tests for multiple comparisons [MO07].

## 3. Results

We first assessed the broad-band oscillatory activity following the presentation of visual motion stimuli. For this, we combined the PRE trials in response to all motion coherence levels for all three experimental groups (*N* = 36), and performed a time-frequency analysis of the MEG responses. A spatio-temporal clustering permutation test corrected for multiple comparisons (see Experimental Procedures) on post-stimulus time-frequency activity (Figure 2.A, left panel) revealed a significant decrease of *α* (alpha: 7 *−* 14*Hz*) power (p *<* 0.001) with a significant increase in the power of broadband high *γ* (60*−*120*Hz*, p *<* 0.001) as compared to baseline.

**Figure 2:**
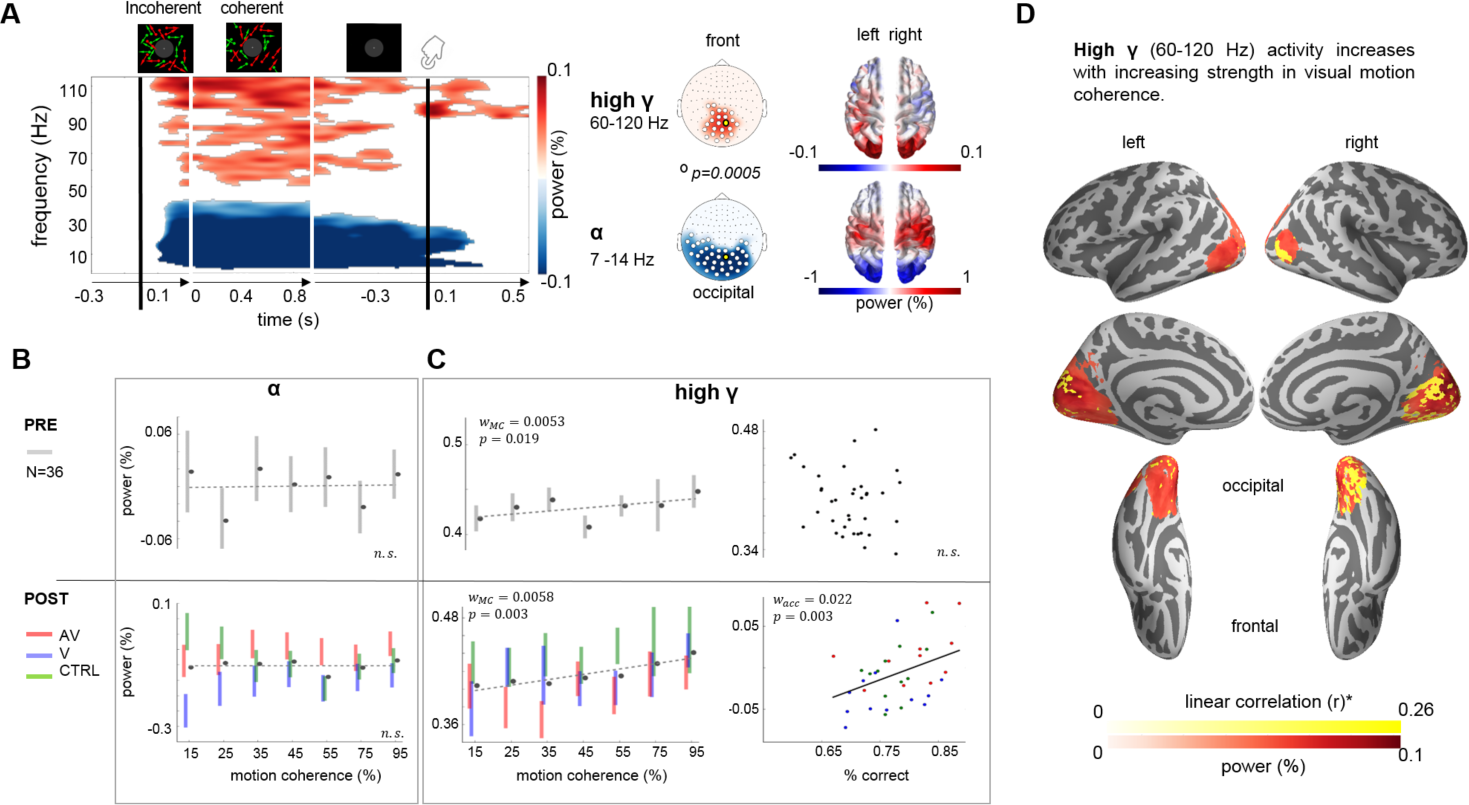
Occipital low-frequency suppression and motion strength (and POST-accuracy) dependent broadband. *γ* **increase**. (**A**) Significant occipital time-frequency clusters of low-frequency (*<* 45 Hz) power suppression and *γ* (45 *−* 120 Hz) band increase were found during the presentation of motion coherence (left panel). Time-frequency analysis was locked to the onset of coherent motion (first black vertical line) and to coherence onset (first white vertical line; the second white vertical is the offset) as well as response-locked (second black vertical line). The three separate analyses stacked together provide the full unfolding of oscillatory activity during the trial. The group average (*N* = 36) time-frequency response of the PRE trials showing a sustained decrease of low-frequency power with an increase in broadband *γ* power is reported for an occipital sensor from the obtained spatial clusters (highlighted yellow sensor in the central panel). Source estimates of *α* (7 *−* 14 Hz) power and broadband *γ* revealed the implication of visual and parietal cortices (right panel). (**B**) In occipital sensors, we found no significant modulations of *α* power as a function of motion coherence in PRE (top panel) or in POST (bottom panel). Bars are 1 s.e.m. (**C**) High *γ* activity increased with motion coherence in PRE (left top panel) and in POST (left bottom), but increased with accuracy only in POST (right bottom). Bars are 1 s.e.m. (**D**) Source estimates showed a significant linear relationship between high *γ* and motion coherence in occipital cortices, supporting the patterns seen at the scalp level.

Significant changes in *β* (15 *−* 30*Hz*) were also found. The significant clusters observed for both the sustained decrease in *α* power and the increase in high-frequency *γ* power were localized in the occipital sensors (Figure 2.A, middle panel). This pattern lasted throughout the presentation of visual motion coherence. Consistent with the topographical pattern at the scalp level, source estimations of the *α* and the high *γ* responses suggested generators located in bilateral visual cortices (Figure 2.A, right panel). This time-frequency pattern during unisensory visual motion coherence was consistent with previously reported time-frequency responses induced by visual motion stimuli [Don+07; Sie+06]. The significant increase in *γ* band during visual motion coherence was also consistent with a previous report of visual motion eliciting a stronger *γ* response than stationary visual stimuli [Swe+09]. We then asked whether the post-stimulus power changes in *α*, *γ*, and *β* were linked to the strength of visual motion coherence in PRE (for all participants) and in POST (as a function of the experimental group), and then proceeded with the exploration of the *β* band.

### 3.1 Occipital α suppression is independent of sensory evidence and training history

In PRE, *i.e.*, prior to any training, we used the grand average data (*N* = 36) and assessed the occipital *α* power from the onset of motion coherence on (Figure 2.A, left panel, white demarcation lines) as a function of motion coherence (Figure 2.B) using non-parametric statistics and a general linear model. We found no significant relationships between *α* power and motion coherence. The same regression analysis was performed in the POST data, independently for each experimental group (*N* = 12) in order to preserve the distinct training history of each group. Again, we found no significant relationships between *α* power and motion coherence, and no significant differences in *α* power between PRE and POST experimental blocks. Overall, we found no substantial evidence that *α* power varied as a function of motion coherence strength or training history. While the absence of systematic occipital *α* modulation limits the functional specificity of *α* in this task, the general decrease of *α* power during stimulus presentation was generally consistent with the inhibitory gating of visual information [JM10; Zum+14] thereby an *α* power decrease could be taken as an index of selective attention [FS11].

### 3.2 Broadband high-γ power increases with the strength in motion coherence and post-training performance

Before training (PRE, *N* = 36), the occipital broadband high *γ* following the presentation of visual motion coherence showed a significant increase with the strength of visual motion coherence (*w_MC_* = 0.0053*,p <* 0.05; Figure 2.C, left top panel). A similar analysis performed on POST data separately for each experimental group (*N* = 12) revealed a significant linear relationship between the post-stimulus *γ* power increase and the increase in stimulus motion coherence. This effect was seen in all three groups irrespective of training history (*w_MC_* = 0.0058, *p <* 0.05; Figure 2.C, left bottom panel). This observation was consistent with the important role of high *γ* power during motion discrimination and its modulation by the strength in visual motion ([Sie+06]. In PRE, no other effects or interactions were found when adding participants’ behavioral correctness (C), reaction times (RT), or confidence ratings (CR) to the general linear model (see Experimental Procedures; additional information regarding behavioral outcomes provided in [Zil+14]). To the contrary, in POST, a positive interaction between correctness and motion coherence drove the regression analysis on its own (*N* = 36, *w_MC−C_* = 0.0055, *p <* 0.005). In fact, irrespective of training history, the interaction between the strength of motion coherence and performance explained the linear relationship between participants’ correctness and occipital broadband high *γ* power (see Figure 2.C, right panels). Subsequent source estimations (see Experimental Procedures) suggested that the increased modulation of high *γ* band power originated in visual cortices (Figure 2.D). This observation is in general agreement with previous findings linking local *γ* band activity to the encoding of sensory evidence [VSS00] during a visual motion discrimination task [Sie+06; Sie+11], and suggested that irrespective of sensory history in training, the reliability of visual sensory evidence contributed to successful task performance.

### 3.3 β power is sensitive to integrated evidence during decision-making

In addition to the post-stimulus *α* and *γ* effects found in PRE (*N* = 36), we also observed two significant *β* clusters (15 *−* 30*Hz*) partially overlapping over the frontal sensors during the presentation of coherent motion: a bilateral early increase in *β* band power (p *<* 0.005) was subsequently followed by a significant decrease (p *<* 0.005) over the left hemispheric sensors. The decrease in *β* band power started around 600 ms following the onset of visual motion coherence (Figure 3.A). The same analysis performed on POST data (*N* = 36) showed, overall, that the decrease in *β* power was left-lateralized and occurred more strongly over left frontal sensors. In what follows, we further investigate changes in *β* power.

**Figure 3:**
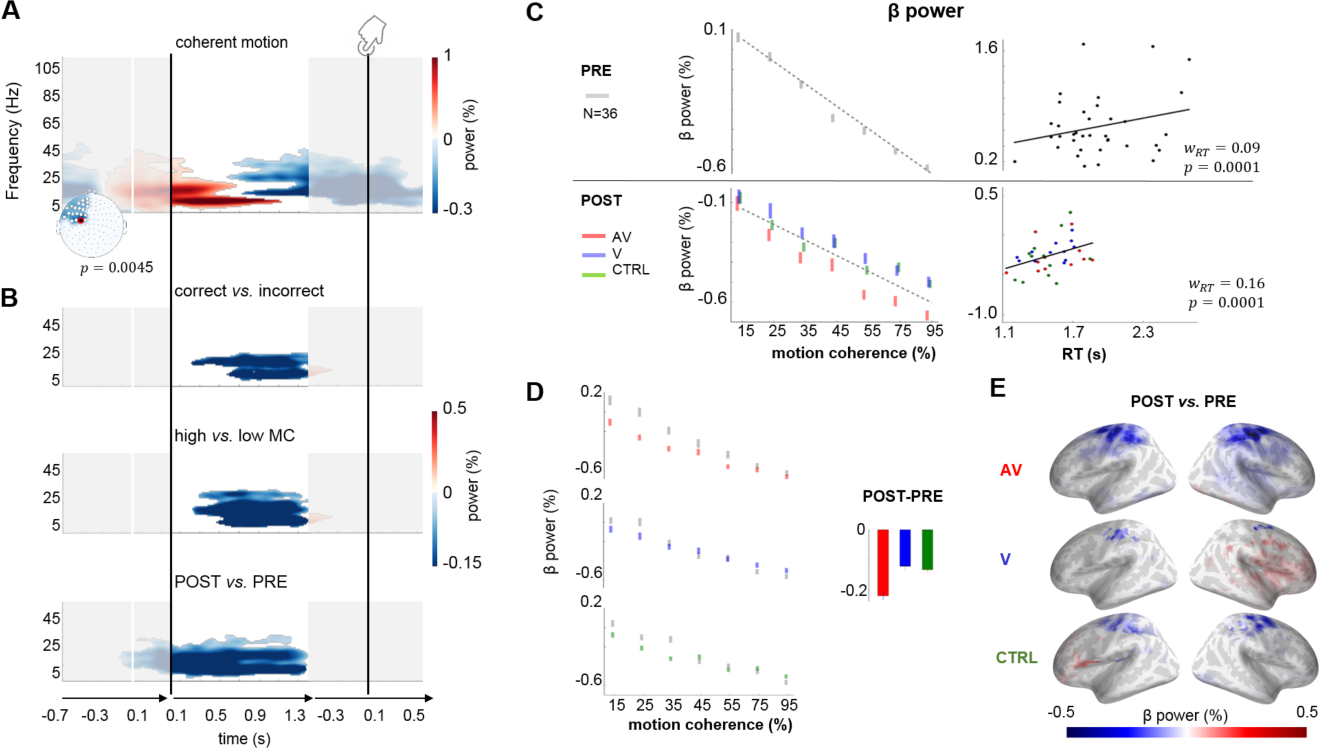
The modulations of *β* power activity indicate a gain in efficiency. (**A**) Group average (*N* = 36) time-frequency maps during the PRE block. The analysis was separately performed on trials locked to the incoherent stimulus onset (white vertical line), locked to the coherent motion (first black vertical line) and locked to the response (second black vertical line). All three analyses were stacked for illustration and provided for a left hemispheric sensor (red dot on left topographic map). Time-frequency permutation clustering statistics revealed two *β* (15 *−* 30 Hz) components partially overlapping over frontal sensors (topography and represented sensor reported on the bottom right corner) during the presentation of the coherent dot motion: a significant bilateral early increase of *β* power (red) was followed by a significant decrease solely over the left hemispheric sensors. (**B**) Statistical contrasts tested the changes in *β* power between correct *vs.* incorrect trials (top panel), high *vs.* low motion coherence trials (middle panel) and POST *vs.* PRE trials (lower panel). All three contrasts revealed a stronger decrease of *β* power. (**C**) Consistent with contrasts in (B), *β* power linearly decreased with increasing motion coherence in PRE and in POST (left top and bottom, respectively) but linearly increased with RT in PRE and POST (right top and bottom, respectively). (**D**) The AV group showed the strongest *β* power decrease from PRE (gray) to POST (red) for all strengths of visual motion coherence (left panel). Bars are 1 s.e.m. Thus, during motion coherence stimuli the strongest overall *β* power decrease from PRE to POST was observed for the AV group (histogram on the right). Source estimates of *β* power showed a significant decrease in POST as compared to PRE over (E) the motor and parietal cortices. This effect was strongest for the AV group.

In a first working hypothesis, we considered prior work showing that changes in *β* power contribute to perceptual decision-making [Don+09; Ala+17], and that motor *β* power can be modulated by attention when anticipating motion coherence onset [Sal+10] and may index functional inhibition during perceptual discrimination tasks engaging different sensory modalities [Cas+01; Bau+12]. To test whether these affected the observed *β* power suppression in our study, we performed cluster permutations on time-frequency data locked to the onset of motion coherence, and devised three contrasts of interest (Figure 3.B): correct *vs.* incorrect trials in PRE as in [Don+09] (top panel), high *vs.* low motion coherence in PRE (middle panel) and PRE *vs.* POST trials (bottom panel). In all three contrasts, we found a significant decrease of *β* power so that the *a priori* easiest trials yielded a larger suppression of *β* power compared to the more difficult trials. Specifically, we found a systematic late decrease of *β* power in the high *vs.* low motion coherence contrast (p *<* 0.01; Figure 3.B, top) and in the correct *vs.* incorrect trials (p *<* 0.05; Figure 3.B, middle). A similar, yet longer-lasting and left-lateralized frontal effect, was found in the POST *vs.* PRE contrast (p *<* 0.01; Figure 3.B, bottom).

As the decrease in *β* power was found locked to the coherence onset but late in the trial – i.e. just before participants’ responses –, it may have reflected the seminal *β* suppression preceding movement onset [PDS99] seen when locking epochs to the individuals’ reaction times (RT) (Figure 3.A, second black line). To test for the possibility that the observed *β* suppression reflected *β* event-related desynchronization shaped by motor readiness and action execution [Mim+99; JB11], we thus locked the trials to participants’ RT and tested the same contrasts as those performed previously on the trials locked to the onset of motion coherence (*i.e.*, correct *vs.* incorrect in PRE data, high *vs.* low motion coherence in PRE data and PRE *vs.* POST). The correct *vs.* incorrect, and the high *vs.* low motion coherence response-locked contrasts did not reveal a decrease but, instead, a small but significant increase (p *<* 0.05) of *β* power before movement preparation. This pattern was only detected when locking the data to the RT (Figure 3.B first two rows on the right) and was distributed over the posterior and frontal sensors. This effect appeared to converge with previous observations [Sie+11], with *β* power suggested to mediate stages of decision-making linking sensory evidence encoding with choice-related action execution. Here, we did not observe significant differences when contrasting the PRE and POST activity for this effect (Figure 3.B third row on the right), and we thus did not pursue the analysis of this specific effect previously investigated in detail. Importantly however, this response-locked *β* activity did not seem to be shaped by sensory history in training and the changes in *β* power locked to motion coherence onset (Figure 3.A and Figure 3.B) were thus distinct from the seminal response-locked effect.

Second, as the modulations of *β* power suppression were not solely specific to the presentation of motion coherence, but also sensitive to the correctness and the type of training participants underwent, we tested whether, in the absence of task, the same *β* power decrease could be seen. For this, we used the localizer data during which participants attended coherent motion stimuli in the absence of a task – namely, during passive viewing. We contrasted brain responses elicited by coherent motion with those obtained in response to incoherent motion. We found no significant *β* power changes in this contrast, suggesting that being engaged in the discrimination task was necessary to observe the *β* suppression effects.

Third, we performed a separate regression analysis (GLM) on the PRE (*N* = 36) and the POST data (independently for each experimental group, *N* = 12) (Figure 3.C, top and bottom panels, respectively). We used the strength of motion coherence and three behavioral variables (correctness, RT, confidence ratings) as regressors. With this approach, we assessed which of the stimulus motion coherence, or of the three behavioral outcomes, contributed most to the variance of the observed modulation in *β* power. We found that *β* power significantly decreased with increasing strength in motion coherence in PRE (*N* = 36, *w* = *−*0.08, *p <* 0.001), (Figure 3.C, left top panel). We also found a significant positive interaction between motion coherence and RT (*N* = 36, *w* = 0.02, *p <* 0.001; (Figure 3.C, right top panel)): In other words, for a given strength of visual motion coherence, we observed a decrease of *β* power with faster RT. Altogether, we thus observed that the strongest visual motion coherence and the fastest RT showed the lowest *β* power.

We then applied the same regression analysis on POST data separately for the three experimental groups (*N* = 12). To make the group-specific results comparable, *β* power from PRE data were separately subtracted from each individual group’s POST data. This analysis revealed a decrease in the slope of the regression between *β* power and the strength of visual motion coherence in all three experimental groups (Figure 3.C, bottom left). This was consistent with the fact that all participants improved their performance after training with increased accuracy, decreased RT, and increased confidence rating [Zil+14]. Similar to the PRE effects, we found a positive interaction between the strength of visual motion coherence and RT with *β* power (Figure 3.C, bottom right). Crucially, an overall decrease of *β* power from PRE to POST was consistently observed for all levels of visual motion coherence in the AV group as compared to the V and the CTRL groups (Figure 3.D). That motion coherence and RT were the main contributors to the *β* power variability was consistent with the observation that all experimental groups were faster in POST as compared to PRE. Interestingly however, that the AV group displayed the largest decrease of *β* power overall after training was also consistent with its overall better performance compared to the other groups (and not with a faster response as the RT were comparable across groups, [Zil+14]). In other words, each group showed an overall decrease of *β* power as a function of the strength of visual motion coherence, which may indicate an overall gain in stimulus processing efficiency as the regression slopes across groups were comparable (Figure 3.D, left panel). Additionally, this decrease was shifted down for the AV group as indicated by the histogram in Figure 3.D, that shows the overall difference (mean and s.e.m. over subjects) of *β* power between POST and PRE for each group, separately. Finally, the performance on the task showed a significant correlation with *β* power but only when using an independent linear regression model, suggesting that motion coherence and RT contributed most to the *β* power effects, which in turn affected performance.

To sum up our observations on *β* power locked to the onset of visual motion coherence, the task-related decrease in *β* power over the frontal sensors got generally stronger with integrated evidence to perform the task. Additionally, congruent multisensory training (AV) induced a larger (POST-PRE) decrease of *β* power than other unisensory (V) or conflicting audiovisual (CTRL) trainings. Consistent with the scalp data, the POST *vs.* PRE statistical contrasts of source estimates showed a strong *β* power decrease over parieto-central regions especially for the AV group; this decrease was also observed in the V and in the CTRL groups to a smaller extent (Figure 3.E). The observed *β* power suppression during motion coherence discrimination converges with previous literature reporting a central role of *β* power during perceptual decision making tasks [Wya+12; Don+09; Ala+17].

As interim summary for the univariate oscillatory analysis, we observed that *α* and broadband *γ* responses during the presentation of visual coherent motion were not significantly affected by training history, in contrast to *β* oscillatory activity seemingly affected by the degree of integrated evidence during training. *β* oscillations may play an important role in (multi)sensory perceptual discrimination consistent with its role in mediating interactions across distant structures during perceptual decision-making [Sie+11]. To disentangle the possible networks mediating these effects, we turned to multivariate functional connectivity (FC) analysis and investigated whether medium- and long-range interactions between cortical regions could provide complementary insights on the specificity of oscillatory regimes as a function of sensory history in training (Figure 1.E).

### 3.4 Task-related network synchronization during visual coherent motion discrimination

To characterize the functional connectivity (FC) induced by (multi)sensory training in the different oscillatory regimes, we estimated the PRE and POST activity during the presentation of motion coherence (i.e., excluding the initial incoherence interval of the stimuli) using 28 cortical regions (ROIs; Figure 4.A). Hence, the bivariate FC was estimated using the weighted phase-lag index (wPLI) (Figure 4.A) in three main synchronization regimes (*α*, *β* and high *γ*). ROIs were selected in a manner orthogonal to the contrasts of interest, mainly by performing a source estimation of the grand average data across all experimental conditions ([Zil+14], see Methods). All statistical contrasts (Figure 4.B) were based on non-parametric permutation t-tests. Only phase coupling values showing significant differences (*p <* 0.01) were retained in the resulting functional networks herein reported.

**Figure 4:**
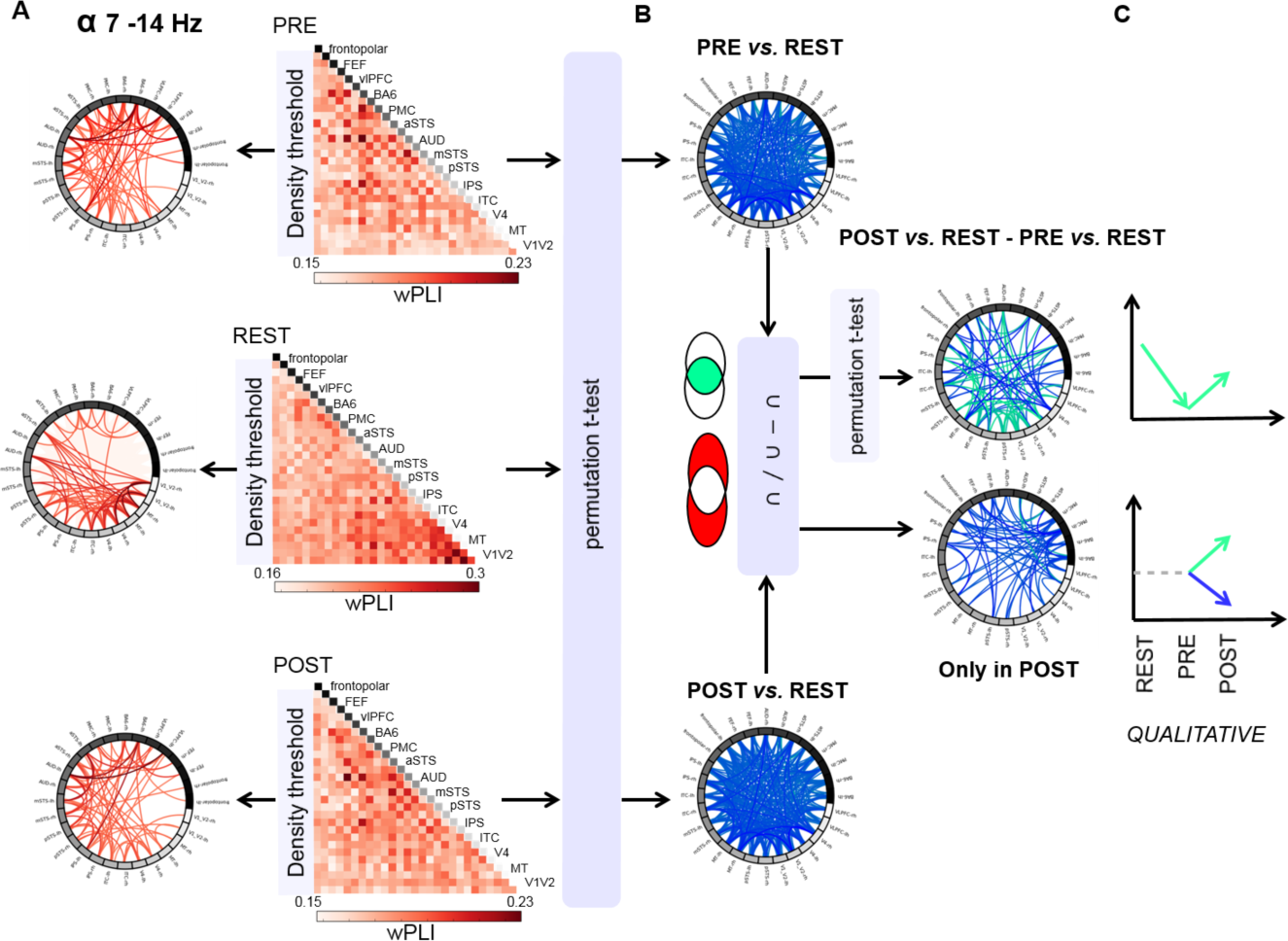
Overview of statistical contrasts performed to extract functional oscillatory networks. (**A**) Functional connectivity (FC) estimates during the presentation of coherent motion, in PRE and in POST, contrasted with resting-state FC patterns (REST). For illustration, we report the full and thresholded *α* oscillatory networks separately for PRE, REST and POST (top to bottom panels, respectively). Statistical contrasts were based on non-parametric permutation t-tests and performed on the 28 cortical regions. (**B**) Pairwise phase couplings were contrasted to show significant differences of weighted Phase Lag Index (wPLI) values characterizing the task-related FC network (PRE *vs.* REST and POST *vs.* REST). (**C**) The FC patterns computed in (B) were compared to assess the variability of the task-related FC in PRE and in POST, as well as to characterize the appearance of new connectivity patterns in POST. FP: frontopolar; FEF: frontal eye field; vLPFC: ventro-lateral prefrontal cortex; PMC: primary motor cortex; BA6: supplementary motor cortex; IPS: intra-parietal sulcus; ITC: inferior temporal cortex; AUD: auditory cortex; aSTS: anterior superior temporal sulcus; mSTS: middle STS; pSTS: posterior STS; MT: middle temporal visual motion area; V4: visual area 4; V1-V2:primary and secondary visual cortices.

First, we estimated the functional connectivity pattern during PRE and POST (*i.e.* during task) that significantly differed from resting-state (PRE *vs.* REST and POST *vs.* REST; Figure 4.B). The subtraction of the resting-state FC from PRE and POST was used as an equivalent of baseline in univariate analyses, and thus was performed to ensure that we characterized the task-relevant FC in both POST and PRE relative to the resting-state network. A direct comparison of POST *vs.* PRE FC without consideration of the initial resting-state FC could be a confounding factor in the interpretation of the results, and could falsely assign significant changes of FC to training effects, when they may in fact simply result from transitioning from REST to task. We then considered the task-relevant network common to PRE and POST (Figure 4.C, top) and explored the effects of training on the changes of cortical interactions and oscillatory couplings.

Consistent with the occipital decrease in *α* band power observed in the univariate time-frequency analysis, we found a significant uncoupling of the *α* oscillatory network in task (both in PRE and in POST) as compared to REST (Figure 5.A, bottom left panel). A relative increase in synchronization modulated by sensory history (Figure 5.A, left column) was found from PRE to POST, involving a large network comprising occipital, temporal and parietal regions. This relative significant increase in *α* synchronization was observed in the V and in the AV groups, but not in the CTRL group. A similar analysis was performed for the *β* and the *γ* oscillatory regimes and, in contrast to *α*, a strengthening of large-scale coupling in the *β* and *γ* bands were observed between REST and task (PRE, POST) (Figure 5.A, bottom panels). The task-related *β* network present in all groups implicated vlPFC, IPS and MT but showed a significant strengthening from PRE to POST solely in the AV group (5.A, middle column). The significant POST *vs.* PRE increase of task-related FC was also observed for the AV group in the high *γ* between the auditory regions and the pSTS. In sum, all three groups displayed a characteristic desynchronization of the *α* network when engaged in the task but a higher synchronization in POST than in PRE of the *α* network for the AV and V groups. Conversely, an increased synchronization of specific *β* and *γ* networks was found in all three groups from resting-state to PRE, and again to POST, but only in the AV group did we see an increase of *β* and *γ* synchronization post-training.

**Figure 5:**
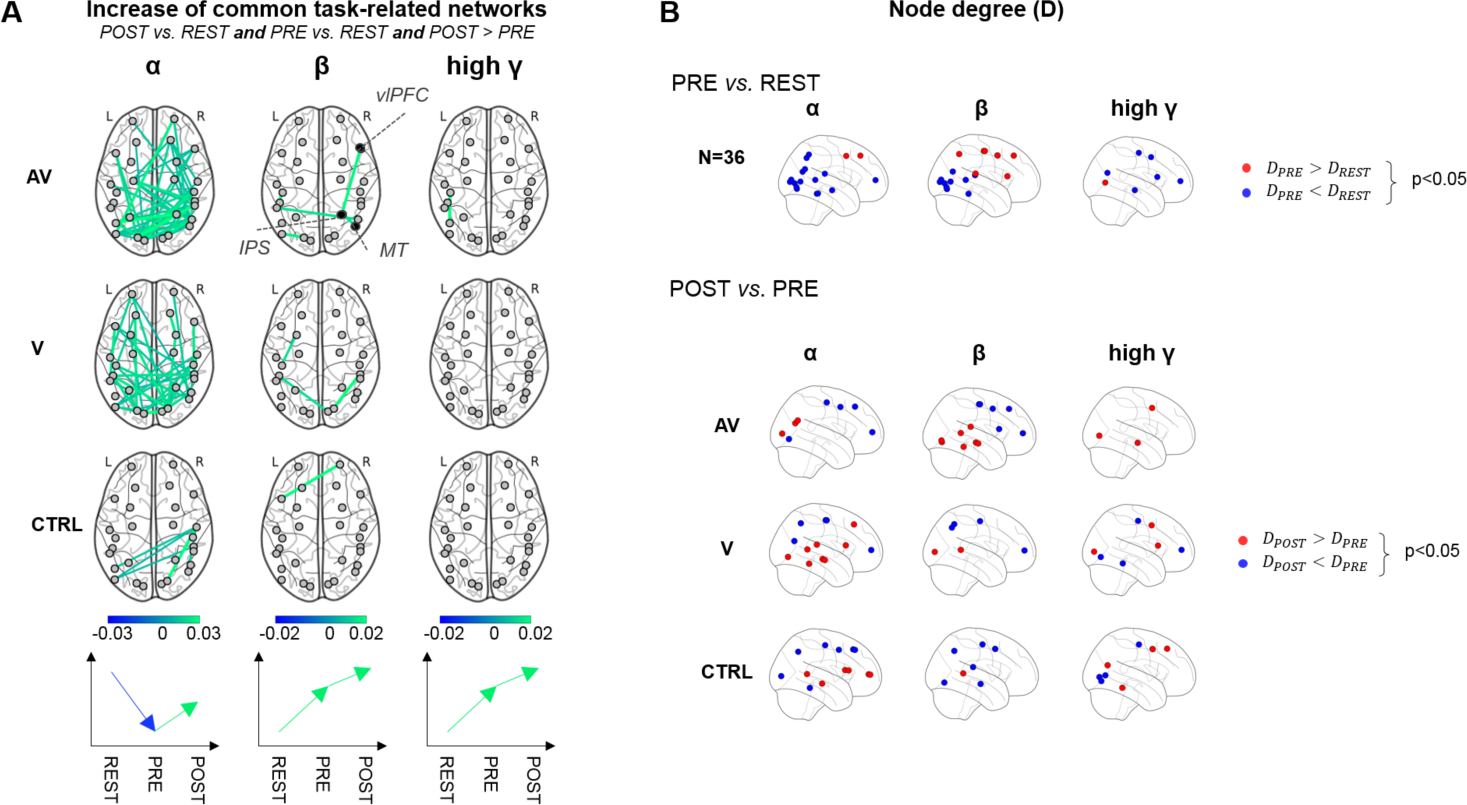
Fluctuations of the task-related networks in pre- and post-training. (**A**) Source estimation was performed to obtain cortical activity in 28 regions of interest (see Section 2.7 for details). Pairwise cortical interactions based on wPLI, and averaged in each frequency band of interest, have been estimated for each condition (REST, PRE and POST). POST vs PRE contrasts of cortical interactions (lines connecting two regions in the figure) within the task-related FC network common to PRE and POST was separately studied for *α* (left), *β* (middle), and *γ* (right) and for the 3 training groups (AV:top, V: middle; CTRL: bottom). A qualitative description of FC changes (significant increases and decreases of interactions) is provided at the bottom, showing that increases from PRE to POST were relative to REST, with an initial desynchronization of *α* from REST to task, and a relative synchronization of *β* and *γ* from REST to task for all three groups. Most notably, both the AV and the V groups showed an overall increase of *α* FC between PRE and POST. Although a significant *β* phase-coupling in task-related FC network linking IPS, vLPFC and MT was found in all groups, only the AV group showed a significant increase of this *β* network following training. (**B**) Topological changes in FC networks from REST to PRE (top) and from PRE to POST (bottom). POST *vs.* PRE contrasts were performed for each training group separately. In the *β* network, the node degree in PRE increased in frontal and parietal regions, whereas it decreased in occipito-temporal regions as compared to REST. The reverse pattern was observed in the *β* network from PRE to POST in the AV group.

### 3.5 Brain network analysis and topological differences in regional connectivity

To investigate the degree of interaction of each brain region, a brain network analysis was carried out using a measure of centrality as index (cf Eq. (3)). This analysis allowed investigating whether specific regions played a central role by assessing the topology of the estimated FC networks based on the number of phase coupling values (connections) over a specific threshold for each ROI (*i.e.* node degree, see Experimental Procedures). This quantification revealed distinct patterns for each oscillatory regime (Figure 5.B), all corroborating our previous analyses (Figure 5.A). The changes in the node degree within the estimated FC networks were assessed with the statistical contrasts PRE *vs.* REST combining all groups, and POST *vs.* PRE on a per group basis.

First, a general task-related decrease of node degree from REST to PRE (Figure 5.B, *α* blue nodes, top left) was observed in parietal, occipital and temporal regions for the *α* oscillatory network. This observation was consistent with the global *α* desynchronization during task as compared to REST. The same contrast for the *β* network (Figure 5.B, *β*, top middle) showed an increase of node degree in PRE in frontal and parietal regions (red nodes), but a decrease in occipito-temporal regions (blue nodes) as compared to REST networks. This pattern was expected considering that long-range cortical interactions in the *β* band are known to involve fronto-parietal regions during perceptual decision-making [Don+07; Sie+11]. Motor cortices also showed a higher node degree in PRE than in REST, reflecting the information flow during task execution mediated by *β* oscillatory networks. The same contrast for the *γ* network (Figure 5.B, *γ*, top right) showed mainly a left-lateralized decrease of node degree but an increase in posterior regions.

We then investigated the changes in FC between POST and PRE as a function of training (Figure 5.B, bottom rows). For the AV group, the analysis of *β* oscillatory networks revealed a clear reversal of the node degree pattern in the POST *vs.* PRE contrast as compared to the PRE *vs.* REST contrast: an increase of node degree from PRE to POST was observed in occipito-temporal regions (mainly in the right hemisphere), while a decrease was found in frontal regions (mainly in the left hemisphere). The node degree value of *β* oscillatory networks implicating motor cortices also decreased with training in all three experimental groups. Conversely, the right mSTS region, which showed a decreasing node degree from REST to PRE, now consistently increased from PRE to POST in all three groups. This suggested the implication of the mSTS during actual training, the synchronization of which got stronger and more extensive (up to visual regions V4) following training.

In the topological analysis of high *γ* oscillatory networks, frontal regions exhibited opposite dynamics as compared to our observation in the *β* band. The node degree of the left frontal BA6 region (pre-motor and supplementary motor regions) decreased from REST to PRE, and increased from PRE to POST for the three groups. These results were in line with previous literature [Don+07] showing that high *γ* and *β* choice-predictive activities showed opposite changes during perceptual decision-making. In the same study [Don+07], oscillatory activities build up gradually during stimulus evidence encoding to reflect the integration of high *γ* activity in MT. Here, on the other hand, the node degree in mSTS and MT regions selectively increased after congruent multisensory training, consistent with the observed selective implication of these regions in the task [Zil+14].

### 3.6 Emergence of β and γ functional networks following training with coherent audiovisual motion

The potential emergence of new functional coupling of cortical brain networks after training (*i.e.*, POST-specific) was addressed on a per group basis (Figure 4.C, bottom). In Figure 6.A, we report cortical interactions that were specific to post-training, *i.e.*, phase couplings among brain regions that were not significantly seen in the PRE *vs.* REST contrast but which significantly emerged after training in the POST *vs* REST contrast (see Methods). These POST-specific couplings between cortical regions emerged in a training-selective manner in the *β* band and only for the AV training group (Figure 6.A). This suggested that multisensory training with temporally comodulated audiovisual stimuli could subsequently affect the organization of cortical interactions during a purely visual discrimination task. The observed reorganization notably encompassed long-range interactions in POST-specific networks around temporal cortices, consistent with the results of the topological analysis shown in Figure 5.B. Additionally, the emergence of high *γ* phase-coupling after training was found mainly in the AV and the V training groups, while POST-specific *α* connectivity emerged in the CTRL group. Intriguingly, the sole behavioral variable relevant to the emergence of the *β* connectivity was participants’ confidence ratings: a significant linear correlation (*N* = 12, *r* = 0.72, *p* = 0.011) was observed so that an increase in POST-specific *β* band connectivity solely observed in the AV group was commensurate with an increase in these participants’ confidence ratings on the task.

**Figure 6:**
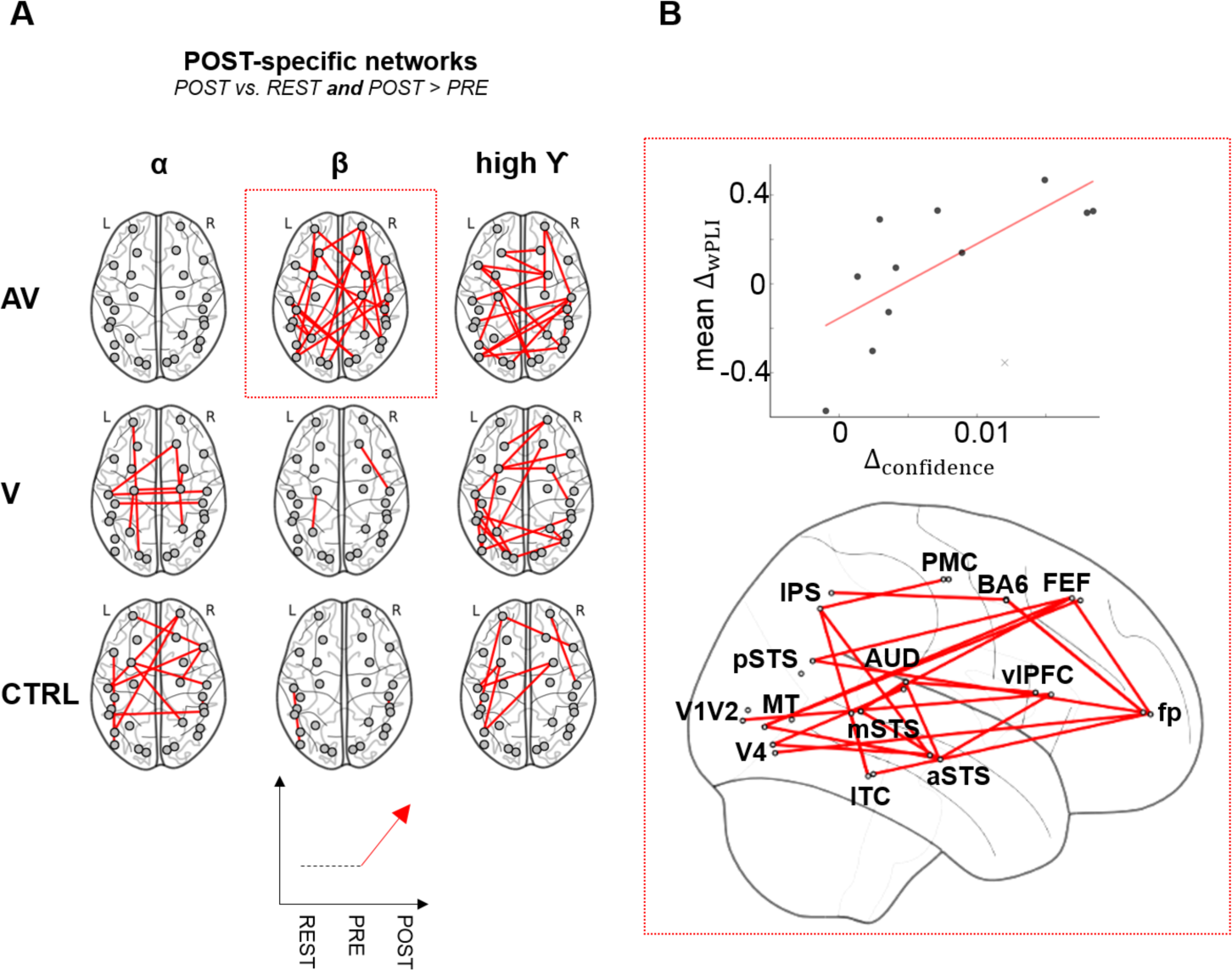
Emerging oscillatory networks following training. (**A**) The phase synchronization based on pairwise wPLI between brain regions (lines connecting two regions in the figure) in *α*, *β*, and *γ* oscillatory regimes was investigated separately as a function of training group. The central finding was the emergence of novel mid- to long-range cortical interactions (POST-specific network, not present during PRE) in the *β* and *γ* networks especially in the AV group. The group trained with incongruent AV stimuli (i.e. CTRL) showed the largest synchronization in *α*. (**B**) A linear correlation between the average increase of the *β* band POST-specific interactions from PRE to POST and the increase in participants’ confidence ratings was observed solely for the AV group (top panel). The *β* band POST-specific network was mainly characterized by fronto-occipital and temporo-parietal interactions (bottom panel).

## 4. Discussion

In this study, we asked how internalized content established on the basis of temporally coherent audiovisual signals could subsequently benefit the discrimination of visual motion coherence. The spectral signatures during the presentation of visual motion coherence included a decrease in occipital *α* and frontal *β* power, and an increase of occipital *γ* power. While the occipital *γ* correlated with the strength in visual motion coherence and post-training performance, *α* activity showed no functional modulation as a function of stimulus property, sensory evidence, performance or training. Additionally, several contrasts revealed that the local *β* power captured an integrated aspect of evidence based decision-making as a function of training history. Second, multivariate functional connectivity analysis based on oscillatory phase coupling showed a relative global increase of *α* (8 *−* 14 Hz) phase synchronization post-training in the V and AV groups as compared to the CTRL group. Third, and importantly, we report the emergence of long-range *β* (15 *−* 30 Hz) and *γ* (60 *−* 120 Hz) synchronization networks implicating temporal, prefrontal, parietal and visual cortices. The emergence of the *β* and *γ* networks was essentially observed following congruent (but not incongruent) multisensory training and the *β* network was indicative of participants’ confidence rating in post-training. Despite the limit represented by the number of participants in this study (*N* = 36), altogether, our results suggest that sensory history in training can subsequently strengthen decision-making networks through the regulation of large-scale oscillatory synchronizations. It would thus be beneficial in the future to increase the number of participants for robust estimation of network changes and characterization. It would also be interesting to test whether similar pattern of beta network can be seen in sensory-impaired populations. The patterns observed in the present study arose from a selected number of cortical regions of interest investigated, which represent a second limit to overcome in future research.

### 4.1 Interplay between top-down α and feedforward γ

Attending stimuli increases both the local and large-scale synchronization of rhythmic neuronal activity in the *γ* band [Eng+01; Fri+01; Var+01; Wan10; Fri15]. An increase *γ* power has been reported during binding ([SG95; Fri+97]), multisensory integration [Bha+02; Mis+07] and semantic congruence across sensory modalities [YGD07; Sch+08a]. Here, we observed an increase in occipital high *γ* band during the presentation of visual motion which, consistent with previous work [Sok+99; Sie+06; Sie+11], increased with increasing strength in visual motion coherence. This effect was seen in visual cortices for all three experimental groups, both before and after their respective training. This pattern converged with the notion that *γ* power provides a spectral index of sensory evidence encoding [Sie+06; Sie+11], and here, may further be a significant indicator of participants’ correct perceptual discrimination following training. As expected [Fri+01; Fri09; Sie+12; JM10], we also observed a concomitant decrease in occipital *α* power. Seminal work has suggested that *α* suppression was stronger for the detection of meaningful objects than for scrambled ones [Van+97], and associated with visuo-spatial [Sau+05] and object-based [FS11] selective attention. An increase in pre-stimulus *α* power is known to impair detection [Bus+09; VD+08; Han+07; Hae+11; Jon+10; ZD10; Gra+17] and, conversely, an increase in *α* power is often observed in unattended modalities [BV10; Kel+06; Wor+00]. *α* oscillations are deemed instrumental for selective attention and the top-down control of information ([SK16]). The modulation of occipital *α* was previously shown to correlate with behavioral improvements of visual motion discrimination in presence of congruently moving sounds [GK14a]. Here, no systematic changes in the occipital *α* were found as a function of experimental or behavioral variables, and this was likely due to differences in paradigmatic and methodological approaches: the most notable one being that we did not directly contrast unisensory *vs* multisensory stimulations *per se*. Rather, the stable level of occipital *α* power over experimental conditions parsimoniously indicated that all participants were effectively attentive to the stimuli irrespective of the strength of motion coherence or training history. Recent work has also suggested that *γ* and *α* (and *β*) oscillations were markers of feedforward and feedback propagation, respectively [VK+14; Wan10; Lee+13; Bas+12; Bas+15; Buf+11; Mic+16; Ric+17]. Given the pattern of stable *α* suppression and increased *γ* power as a function of motion coherence (and post-training correctness), one possible working hypothesis is that, given a stable and sustained endogenous attentional control exerted in the *α* range, perceptual training may improve the efficiency with which *γ* activity propagates sensory evidence up the hierarchy.

### 4.2 The effect of sensory history in training on α and γ network phase-synchronization

In line with the notion that *α* oscillations actively contribute to the selection of cortical regions during task [PP11; PP07], an increase of global *α* phase synchronization was observed from resting-state to task in all groups. Yet, and perhaps more importantly, a global *α* synchronization in parieto-frontal network further increased post-training in the control group trained with distracting or uncorrelated sounds and, to some extent, in the V group – but not the AV group. The presence of large-scale *α* synchronization in the CTRL group could be primarily explained by the selective function of *α* networks, which may help decouple brain regions during conflicting inputs (CTRL). At the same time, while the type of sensory history in training did not selectively affect occipital *γ*, we observed a global strengthening of the post-training *γ* synchronization network. *γ*-band synchronization in brain networks is fundamental in cortical communication as phase-coupling across brain regions may promote the transmission of information across large-scale neuronal networks [Fri09; Fri15; Bas+15]. Global *γ* synchronization is notably considered to denote “*effective, precise and selective*” communication [Fri15]). In this study, training may have improved long-range *γ* synchronization with a possible gain in communication efficiency consistent with the general improvement on the task observed in all groups [Zil+14]. Sensory history in training affected the general increase of *γ* phase synchronization so that the group with the largest behavioral improvement, *i.e.* the AV group, also showed the strongest increase followed by the V and the CTRL group. Altogether, theses results suggest that the type of sensory inputs during a very short training (here a total of 20 minutes) can selectivity affect the coordination of brain regions implicated in the endogenous control of information processing.

### 4.3 β oscillations as integrated evidence

Very recently, *β* oscillations have been proposed to be markers of internal content [SH17] and supramodal processing [Hae+17]: following learning, rhythms such as *β* oscillations may regulate the feedback processing of sensory analysis [BM16] based on abstract categorical representations. These internal network dynamics were observed in prefrontal cortices, and operated in the *α*/*β* bands [BM16]. An important finding in our study is the fundamental role of *β* oscillations, both as local power decrease during sensory encoding and decision-making in all groups, and as an emergent large-scale network following congruent multisensory AV training. Several studies have reported an increased coherence or synchronization in the *β* band associated with multisensory stimulation [Ste+99; Mer+15] and, consistent with the implication of the *β* band in sensorimotor processing [EF10], *β* activity was related to participants response speed in multisensory context. Gleiss and Kayser [GK14b] reported an early difference in *β* band activity but did not find any correlation with behavior. Consistent with another study [Mer+15], we found that modulations of *β* power were observed when participants were engaged in the task but not during passive viewing, and that RT was the main contributor for this effect: the decrease in local *β* power was found for contrasts in which evidence-based decisions were most successful (*i.e.*, for strongest as compared to weakest visual motion coherence, for correct as compared to incorrect trials and for post-compared to pre-training trials). Under the working hypothesis that abstract internal content [BM16; Wut+18] has been learned to drive the processing of incoming sensory information, the strengthening of the large-scale *β* coupling in the group that has received congruent AV training would further suggest that performing the task with temporally coherent audiovisual events strengthened the ability to predict motion coherence in vision. In other words, the changes in *β* power and network synchronization may capture endogenous top-down activations of task-relevant (supramodal) cortical representations, which facilitate communication between brain regions [Kop+00; Sie+11; SH17; Hae+17; BR15]. In this context, recent predictive coding models drawing from audiovisual speech processing [VW+05; Was13; Arn+11] and neurophysiological work [Ric+17; BR15] have pushed forward the notion that prediction errors from one sensory modality to another may be communicated in the *γ* range, whereas top-down predictions may be mediated by *β* oscillations [Arn+11]. As the emergent *β* network was solely seen for the AV group in which it was linearly related to confidence rating, we speculate that the hypothesized combined effects of increased communication efficiency in a feedfoward *γ* network, the endogenous selectivity in the *α* network, and the predictive *β* propagation in the AV group may all contribute to the local selective changes previously reported in MT as a change in the neurometric threshold [Zil+14]. The notion that internal content (as supramodal or abstract representations) may constrain sensory analysis early on provides additional evidence for the implication of large-scale neural oscillations in integrative and predictive brain functions.

Taken together, our results support the notion that cortical computations encompass sensory-based processing and that, consistent with the role of prefrontal cortices shifting activity from feedforward inputs to internal dynamics [BM16], the internal content shaped by multisensory inputs during training can strengthen the selectivity of large-scale oscillatory networks for later adaptive purposes.

## Acknowledgments

This work was supported by ANR-16-CE33-0020 MultiFracs, France, and the Marie Curie IRG-249222 and the ERC-YStG-263584 to V.vW. We thank Dr Laetitia Grabot, Dr Sophie Herbst, and Dr Tadeusz Kononowicz for their comments on the initial version of the MS.

## Footnotes

1 http://surfer.nmr.mgh.harvard.edu/

2 https://github.com/mne-tools/mne-python/blob/master/tutorials/plot_artifacts_correction_ica.py\

3 https://surfer.nmr.mgh.harvard.edu/fswiki/CorticalParcellation

